# PIP2 Influences the Conformational Dynamics of Membrane bound KRAS4b

**DOI:** 10.1101/608265

**Authors:** Mark A. McLean, Andrew G. Stephen, Stephen G. Sligar

## Abstract

KRAS4b is a small GTPase involved in cellular signaling through receptor tyrosine kinases. Activation of KRAS4b is achieved through the interaction with nucleotide exchange factors while inactivation is regulated by through interaction with GTPase activating proteins. The activation of KRAS4b only occurs after recruitment of the regulatory proteins to the plasma membrane thus making the role of the phospholipid bilayer an integral part of the activation mechanism. The phospholipids, primarily with anionic head groups, interact with both the membrane anchoring hypervariable region and the G-domain, thus influencing the orientation of KRAS at the membrane surface. The orientation of the G-domain at the membrane surface is believed to play a role in the regulation of KRAS activation. Much of the research has focused on the role of phosphatidyl serine but little has been done regarding the important signaling lipid phosphatidylinositol-4,5-bisphosphate (PIP2). We report here the use of fluorescence anisotropy decay, atomic force microscopy, and molecular dynamic simulations to show that the presence of PIP2 in the bilayer promotes the interaction of the G-domain with the bilayer surface. The stability of these interactions significantly alters the dynamics of KRAS4b bound to the membrane indicating a potential role for PIP2 in the regulation of KRAS4b activity.

---

RAS proteins are small GTPases that function as molecular switches. Growth factor engagement with extracellular receptors recruits guanine nucleotide exchange factors (GEFs) to the inner leaflet of the plasma membrane (PM) which convert inactive RAS-GDP to active RAS-GTP. Once in the activated state RAS interacts with its primary effectors, RAF kinase, PI3K and RALGDS. Recruitment of these effectors to the PM by RAS results in their activation and the initiation of signal transduction, thereby promoting cell proliferation, differentiation and growth^1^. GTP hydrolysis, catalyzed by GTPase activating proteins (GAPs), results in the inactivation of RAS and termination of growth factor signaling.

RAS mutations have been identified in 20% of human cancers^2^. KRAS is responsible for nearly 85% of all the RAS driven cancers and found with high prevalence in pancreatic, colorectal and lung cancers^3^. The most frequent oncogenic mutations in RAS are localized at positions 12, 13, or 61. These mutations impair GAP stimulated GTPase activity rendering RAS locked in the GTP bound active state, uncoupled from growth factor stimulation.

RAS proteins consists of a G-domain (residues 1 – 165) that binds guanosine nucleotides and a hypervariable region (HVR, residues 166-188) that is involved in anchoring RAS to the membrane. The G-domain can be described as a bi-lobal structure. Lobe 1 consists of residues 1 - 86 while lobe 2 consists of residues 87 – 166. RAS proteins also contain a CAAX recognition sequence at the C-terminus that targets the protein for post translational modification to contain a farnesylated, methylated C-terminus at CYS 185. The RAS isoforms HRAS, NRAS and KRAS4a are reversibly palmitoylated in the HVR. This functions as a second signal to target these proteins to the plasma membrane (PM). KRAS4b, however, is not palmitoylated but contains a poly-LYS sequence that interacts with the anionic lipid head groups at the PM^4^.

In recent years many researches have turned their attention to the interaction of RAS and its effectors with the lipid bilayer. Since all signaling through RAS must occur at the membrane surface, understanding the details of how the bilayer helps control or assemble the signaling complexes may lead to new druggable targets. Abankawa and coworkers showed that the orientation of the G-domain is tuned by the HVR and α-helix 4 which in turn effects recruitment of effectors.^5^ KRAS4b-GDP and KRAS4b-GTP on Nanodiscs containing 20% DOPS exist in different orientations as demonstrated by NMR paramagnetic relaxation enhancement experiments.^6^ Surprisingly, KRAS4b-GTP bound preferentially in an orientation incompatible with effector binding, whereas KRAS4b-GDP resides in an exposed conformation available to effector recognition. Interestingly, the G12D oncogenic mutation and Rasopathy associated mutants K5N and D153V altered the orientation from the occluded to the exposed orientation.^7^ Molecular Dynamics (MD) simulations of KRAS4b-G12D-GTP on bilayers containing 20% POPS identified two membrane bound orientations: OS1 with the effector binding site exposed and OS2 with effector binding site occluded.^8^ A recent study that included a 20 μs MD simulation and single molecule FRET measurements identified these two major orientations.^9^ In this work a third population was observed, that represents an intermediate and possibly a transition state between OS1 and OS2, termed OS0. The orientations seen in the simulations correlated well to the distribution of distances measured by single molecule FRET experiments.

The major focus of many experimental and computational studies has been the role anionic lipids, specifically phosphatidyl serine (PS), play in KRAS membrane orientation. In a recent review^10^ (and references within) the specificity of the KRAS for phospholipids is emphasized by the segregation and nanoclustering of RAS in PS rich environments. This specificity can be altered by single amino acid substitutions in the HVR region.^4^ There has been significantly less effort focused on the role phosphatidyl inositides play in controlling membrane KRAS interactions. While PIP2 has not so far been directly implicated in KRAS activation, a key downstream effector is PI3K whose substrate is PIP2. In addition, the GEF Son of Sevenless (SOS) is known to bind phosphatidyl inositides.^11^ We and others have recently shown that PIP2 interacts with RAS HVR as well as the RAS G-domain and influences the interaction of KRAS4b with the membrane by forming bifurcated salt bridges with positively charged amino acids.^12–14^ In this report we focus on the role of PIP2 in defining the orientation states of KRAS4b using a combination of fluorescence anisotropy decay, atomic force microscopy and long time molecular dynamics simulations.

## MATERIALS AND METHODS

### Nanodisc Preparation

Membrane Scaffold Proteins (MSPs) were expressed and purified and the 6xHIS tag cleaved as previously described.^15^ Ru(bpy)_2_(5-iodoacetamido-1,10-phenanthroline) (RubpyIA) was a kind gift of Dr. Lionel Cheruzel. MSP D73C was labeled at position 73 with RubpyIA by addition of a 4-fold excess TCEP to reduce the free sulfhydyls. 50 mM RubpyIA dissolved in dry DMSO was added to the reaction to achieve a 10:1 RubpyIA:MSP ratio. The reaction was allowed to proceed in the dark for 4 hours with gentle stirring. DTT was added to a final concentration of 2 mM to quench the reaction. Excess dye was removed using a G-25 column equilibrated in 20 mM HEPES pH 7.3, 150 mM NaCl. Labeling efficiency was calculated using the extinction coefficients for MSP ε_280_ = 18.45 mM^−1^ and RubpyIA ε_454_ = 16.6 mM^−1^ ε_280_ = 64.5 mM^−1^.^16^ Nanodiscs were prepared as previously described.^15^

### Fluorescence Anisotropy Measurements

Fully processed KRAS4b-GDP (KRAS-FME) was expressed and purified as previously described.^17^ Samples contained ~100 nM RubpyIA labelled Nanodiscs, 7 μM KRAS-FME, 20 mM HEPES, 150 mM NaCl, and 1 mM MgCl_2_. Fluorescence lifetime and anisotropy decay measurements were performed using a ISS inc. K2 fluorimeter equipped with the K520 Digital Frequency Domain using laser excitation at 486 nm. The frequency dependent phase shift and modulation ratio were fit to a biexponential anisotropy decay according to the methods reviewed by Lakowicz using either the Vinci (ISS Inc.) software package or in house Matlab routines^18^. The intensity decay model used includes a single fluorescence lifetime of ~ 900 ns and two rotational correlations times (θ_r_). Our model consists of a fast rotation ~ 10 ns corresponding to fast fluctuations of the dye molecule and a slower rotation >100 ns corresponding to Nanodisc rotational diffusion. Due to the long lifetime of dye, the fast rotational correlation time is not well resolved but is needed to account for the limiting anisotropy of 0.18, which is a fixed parameter.^16^ KRAS4b binding experiments were carried out at a single frequency as described by Gregory *et al.*^12^

### Atomic Force Microscopy

Atomic force microscopy (AFM) was performed using a Cypher ES Atomic Force Microscope (Asylum Research) running in contact mode in buffer containing 20 mM HEPES, 150 mM NaCl, and 1 mM MgCl_2_ at pH 7.3. Nanodiscs were adsorbed to freshly cleaved mica followed by a buffer rinse and incubation with KRAS4b-FME. The spring constant of the AFM probes were measured prior to each use. Trace and retrace images were then taken at a constant force of 200 pN.

### Molecular Dynamic Simulations

All Molecular dynamics inputs were generated using CHARMM-GUI membrane builder and the CHAMRMM36m forcefield. Production runs were performed using the GROMACS software package.^19–21^ The KRAS4b structure was based on pdb 4OBE with the missing helical HVR region added.^22^ The resulting structure was nearly identical to the structure of full length KRAS4b bound to PDEδ (PDB 5TAR).^23^ Due to the known flexibility of the HVR region the starting structures were generated by five annealing simulations to relax the HVR structure (see supplementary figure S1). Molecular dynamics simulations of planar bilayer systems were performed using the Highly Mobile Membrane Mimetic containing DMPC and 10% DMPS or 10% DMPS / 3 % PIP2.^19,24^ The systems contained 240 to 300 phospholipids, 150 mM KCl resulting in systems that contained 100000 – 110000 atoms. Production runs were performed with an NPT ensemble at 303 K for 1 μs at each of the 5 starting configuration and each lipid composition, for a total of 10 μs. Nanodisc simulation inputs were generated by the CHARMM-GUI Nanodisc builder and production runs were performed on an NPT ensemble at 278 K. Visualization and analysis of the trajectories were performed using VMD and in house Matlab routines.^25^

### MD Analysis

MD trajectories containing frames every 500 ps were analyzed by computing the contacts between KRAS4b and the lipid headgroups defined as any heavy protein atom within 3.2 Å of a head group heavy atom. Residence times lipid-protein contacts were calculated for each KRAS4b residue that comes into contact with a headgroup and counting the number of consecutive frames they remain in contact. The mean residence time is the average of all contacts over the course of all simulations (10 μs total). In addition we employ the definition of G-domain orientation described by Prakash *et al.* ^9^ Briefly, the angle of orientation is defined as the angle, θ, between the bilayer normal and a vector running the alpha carbon of residue 5 and the α-carbon of residue 9. The distance is defined as the difference in the z component of the coordinates for the α-carbon at residue 132 (α helix 4) and the α-carbon at residue 183 which reside just above the buried farnesyl group at residue 185, thus defining distance to the bilayer surface (see supplementary Figure S2). The distribution of the orientation is then visualized using a two dimensional histogram of cos(θ) vs distance.

Finally we define a new parameter, orientational mobility, to analyze the trajectories. For each time step of 500 ps we calculate the angle and distance to the bilayer as described above. Mobility is calculated as the distance between consecutive points on a plot of cos(θ) *vs.* distance during a trajectory and represents how fast the orientation is changing over the time step of the simulation. This parameter can then be analyzed as function of time, distance or orientation angle. For a detailed description of mobility see supplemental Figure S2.

## RESULTS AND DISCUSSION

### Binding of KRAS4b to Anionic Lipids

We performed multi-frequency anisotropy decay measurements of GDP bound KRAS4b interacting with Nanodiscs (ND) containing DMPC with 30% DMPS, or DMPC with 10% PIP2. This labeling scheme differs significantly from our original report as the dye is now tethered to a mutant of the MSP, Cys 73, where as previously we labeled free amino groups on the protein^12^. The ruthenium dye is now site specifically attached to the Nanodisc which removes the complication of multiple labeling sites on the MSP belt. The anisotropy decay is a measure of the rotational diffusion of the KRAS/ND complex. Figure 1 shows a plot of Delta-Phase and Modulation Ratio *vs*. frequency of 30% DMPS Nanodisc (top panel) and 10% PIP2 Nanodiscs (bottom panel) with KRAS4b bound (blue) and without (red). A clear shift in delta phase and modulation ratio indicates that the apparent rotational diffusion rates increase. Due to the tight interactions of α-helix 4 with PIP2^12^, our initial hypothesis to describe the rotational motion of KRAS bound Nanodiscs was that discs in which the G-domain is loosely associated or standing up would display a *slower* rotational correlation time while a complex with the G-domain tightly associated with the bilayer would be more compact and display a *faster* rotational correlation time. Table 1 summarizes the rotational correlation times for KRAS bound to 30% DMPS Nanodiscs and 10% PIP2 Nanodiscs. It is clear that the KRAS Nanodiscs display a *slower* rotational correlation time when they contain 10% PIP2 compared to 30% DMPS.

**Figure 1.**
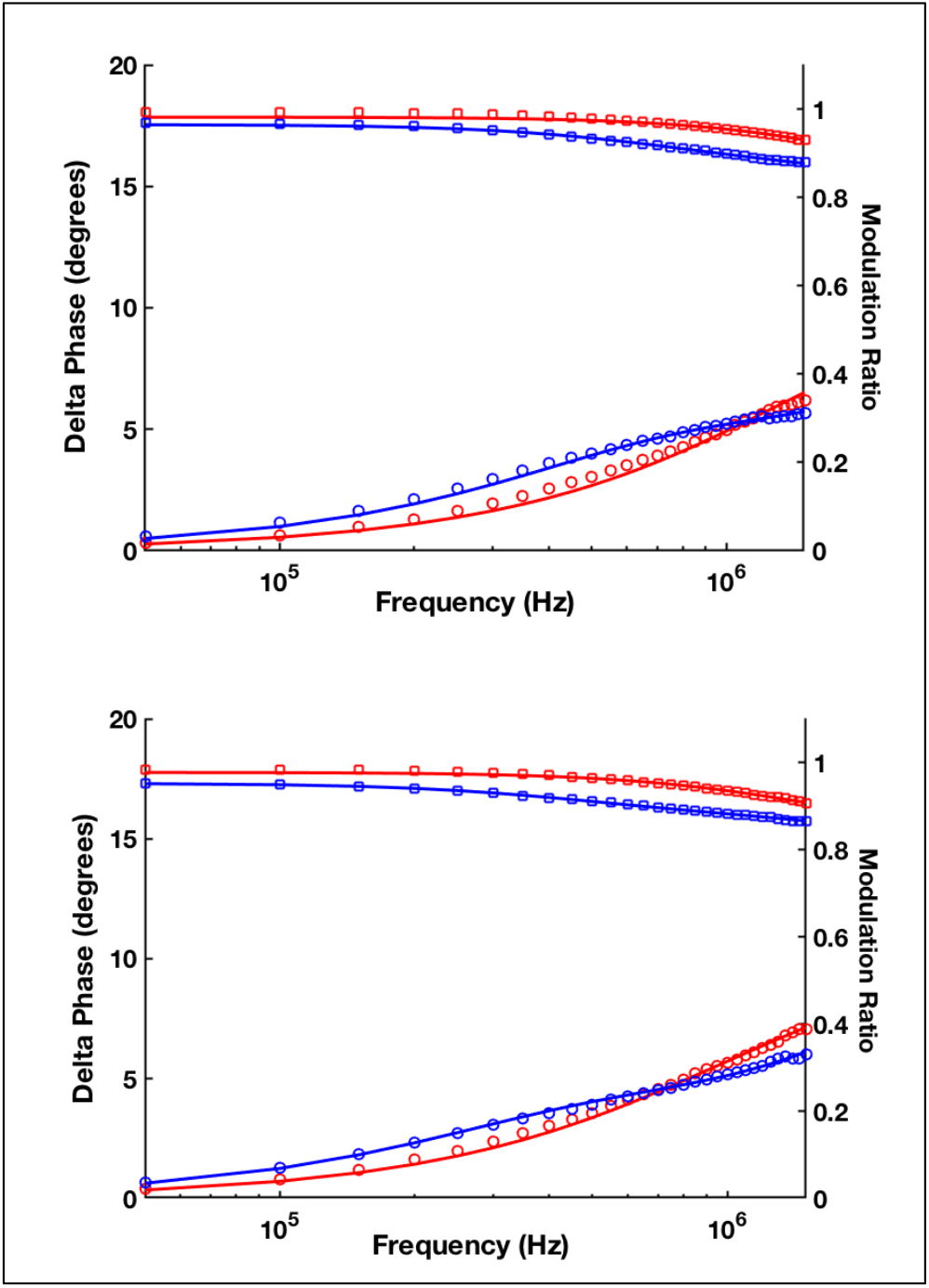
Multifrequency Anisotropy Decay. Top Panel 30% DMPS, Bottom Panel 10% PIP2, Red-Nanodiscs, BlueNanodiscs + KRAS4b. Squares - Modiulation Ratio – Cirgles – Delta Phase. Solid lines represent a fit to two rotational correlation times as described in the methods.

**Table 1.** Rotational Correlation times of KRAS4b Nanodiscs

|  | 30% DMPS | 10% PIP2 |
| --- | --- | --- |
| | $\theta_r$ (ns) | $\theta_r$ (ns) |
| Nanodiscs | $110 \pm 6$ | $102 \pm 9$ |
| Nanodisc-KRAS4b | $258 \pm 10$ | $422 \pm 8$ |

In order to further the understand the contradiction in the observed rotational correlation times and our hypothesis we performed an all atom MD simulation of DMPS Nanodiscs bound to KRAS4b. To estimate the rotational motions of the Nanodisc we calculated the displacement velocity of residue 73 in one of the MSP belts and the radius of gyration of the complex as a function time. The center of mass of the Nanodisc was recentered at each step of the trajectory to remove any translation motions. Therefore, the displacement velocity of residue 73 is a report of the rotational motions of the Nanodisc. The radius of gyration of the Nanodisc-KRAS complex and displacement velocity of the MSP belts were smoothed using a moving box with a width of 50 ns to average out rapid molecular motions and plotted as a function of time. Figure 2, top panel, shows a comparison of the radius of gyration and the MSP displacement velocity. Looking at the radius of gyration (red) we see a sharp decrease as the G-domain comes in contact with the Nanodisc surface. One might expect to see the MSP velocity increase as the radius of gyration increases, but this is clearly not the case. Instead as contacts between the G-domain and lipid head groups form, the MSP velocity decreases (Figure 2, bottom). In our experiments the fluorescent label is attached to the protein belt of the Nanodisc and thus reports on the Nanodisc rotations. Due to the flexible nature of the HVR it appears that the rotational motion of the disc moiety is uncoupled from the motion of the overall complex. Our interpretation is that when the KRAS4b is tethered only by the HVR, there is a significant amount of Nanodisc motion that is uncoupled to that of the entire complex. It is only when the G-domain comes in contact with the disc surface does the rotational motions of the disc correlate with that of the complex. Our previous studies showed that PIP2 influences KRAS4b affinity due to the long lived salt bridges that are formed with both the HVR and the G-domain^26^. Consistent with those findings, our new results suggest that the measured rotational correlation times support a model where the G-domain is more loosely associated with the bilayer surface on PS containing membranes, while on membranes containing PIP2, the G-domain predominantly resides at the Nanodisc surface.

**Figure 2.**
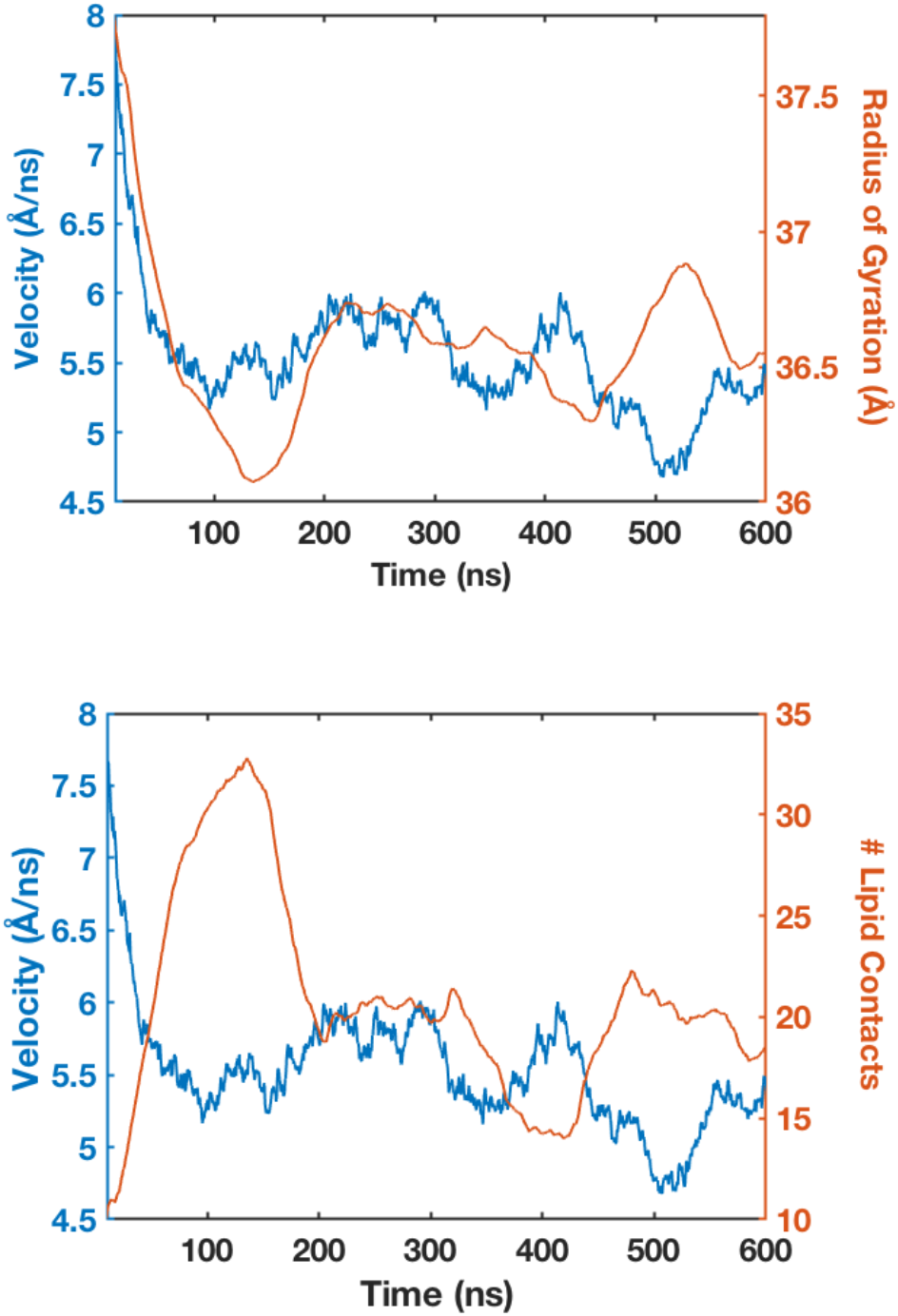
KRAS4b mobility *vs.* time. Top Panel: MSP velocity (blue), radius of gyration (red). Bottom Panel: MSP velocity (blue), Lipid Contacts (red).

### Atomic Force Microscopy of KRAS4b – Nanodiscs

In an attempt to get a structural picture of KRAS4b bound to the Nanodisc surface, we performed contact mode Atomic Force Microscopy (AFM) imaging. Figure 3 shows images of Nanodiscs alone (panels A and B) and Nanodiscs bound to KRAS4b (panels C and D). Nanodisc samples in the absence of KRAS4b appear monodisperse, with the expected diameter of ~10 nM. Addition of KRAS4b to the 30% DMPS sample results in a significant deterioration of image quality (Figure 3C). Discs and some features above the discs can be seen but are not consistent with a tight association of KRAS4b to the bilayer. In stark contrast, the addition of KRAS4b to 10% PIP2 Nanodiscs (Figure 3D) results in a crisp image of Nanodiscs with significant features appearing above the Nanodisc surface. The features, which are presumably KRAS4b, protrude approximate 2 – 3 nm above the disc surface. The diameter of the KRAS4b on the surface appear to be 30 – 50 nm, which arises from a convolution of the AFM tip radius of 20 – 30 nm interacting with a single KRAS4b ^27^.

**Figure 3.**
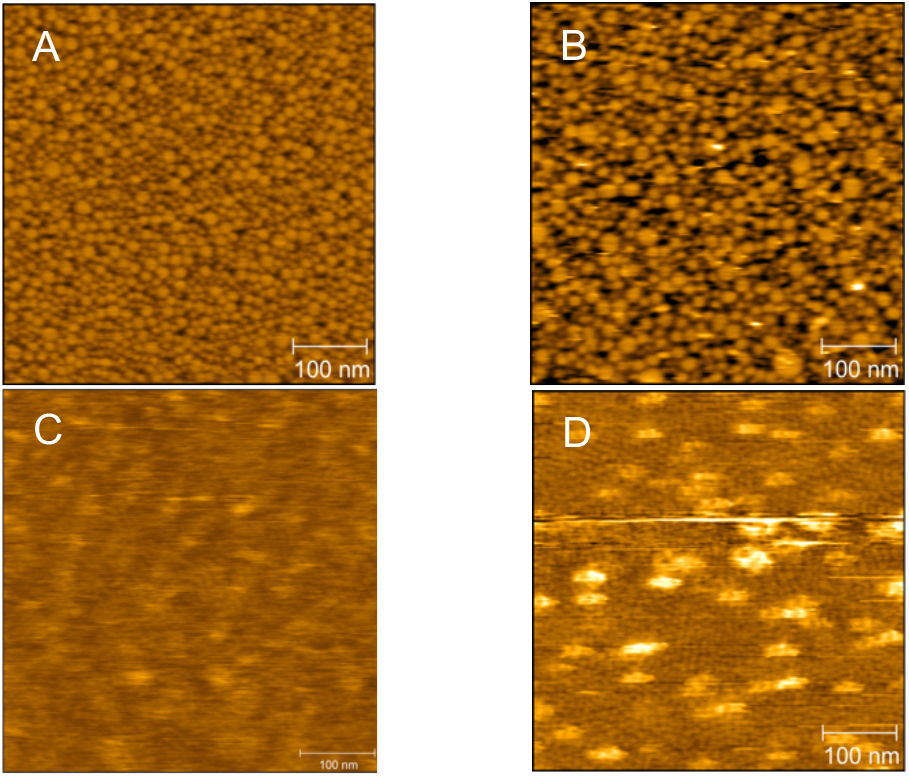
Contact mode AFM images of KRAS4b bound to Nanodiscs. (A) 30% DMPS Nanodiscs. (B) 10% PIP2 Nanodiscs. (C) 30% DMPS Nanodisc-500 nM KRAS4b. (D) 10% PIP2 Nanodiscs-100 nM KRAS4b

The poor image quality of the 30% DMPS-KRAS sample can be attributed to the inherent mobility of KRAS bound to DMPS bilayers. In contact mode AFM imaging, the x and y axis of the image are termed the fast (x) and slow (y) axes. The image is constructed by scanning the AFM tip over all values of x at position y, then moving to a new y position and scanning x again. Each y position is scanned twice, once forward (trace) and then backward (retrace), producing two images one for the trace and another for the retrace. For an ideal hard sample, these two images should be identical. Comparison of these two images thus allows identification of unstable samples or samples that are perturbed by the force of the AFM tip. Here we have leveraged this feature of AFM imaging to gain insight into the mobility of KRAS4b on the Nanodisc surface. Figure 4 shows line traces (height *vs*. x) for several y values from 30% DMPS-KRAS images (left) and 10% PIP2-KRAS (right). The blue lines are extracted from the trace image and the red lines from the retrace image. In the 30% DMPS image we see that the features rarely extend higher than 1.5 nm above the surface. In addition, the trace and retrace profiles do not overlay well. The peaks are consistently shifted in the direction of the trace scan. This is due to the mechanical “pushing” of the KRAS4b in the direction of the scan. This also explains the smaller heights observed on 30% DMPS since the KRAS4b will move out of the way of the tip before the tip reaches the top of the protein. A similar analysis of KRAS4b bound to 10% PIP shows that the line traces consistently overlay with each other and display very little shift between trace the retrace. Moreover, the heights of 2 to 2.5 nm above the Nanodiscs are more consistent with a 20 kDa protein bound to the lipid surface.

**Figure 4.**
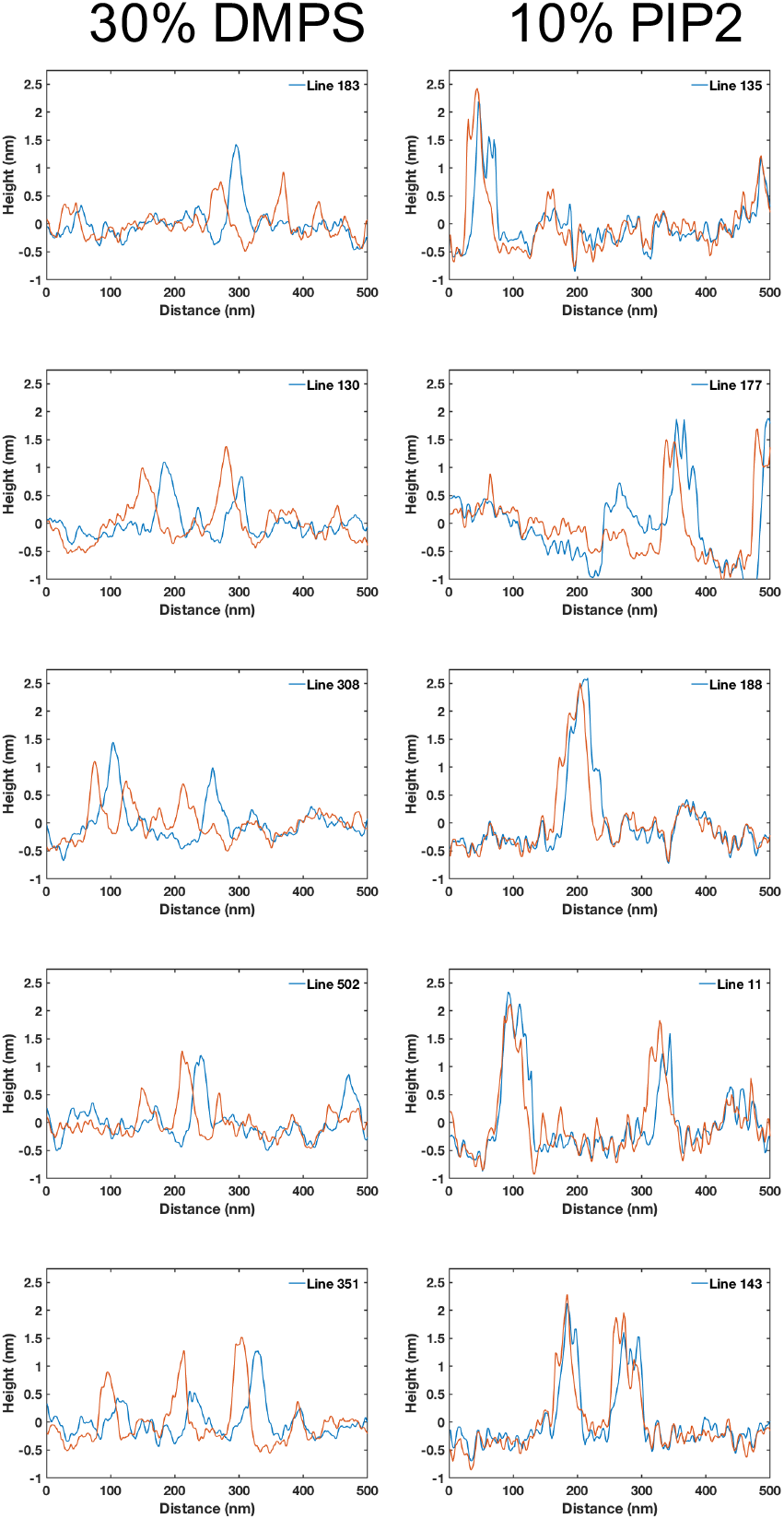
AFM Line traces. Each panel represents a single line of an AFM image. Blue lines are the forward scans and the red lines are the reverse scans.

The AFM imaging results paint a picture of how KRAS4b interacts with DMPS and PIP2 containing bilayers. Our previous findings revealed that KRAS4b binds to 10% PIP2 binds to with an affinity 5 – 6 times tighter than that of 30% DMPS. ^26^ At conditions where the KRAS4b concentration is at ~50% of the dissociation constant, the number of KRAS4b bound to the surface in both images should be similar. The inability to get stable images on 30% DMPS Nanodiscs suggest that KRAS is loosely associated with the bilayer and the G-domain is either standing up and easily pushed aside by the forces of the AFM tip, or associated with the bilayer, as seen in the PIP2 images, but disrupted by the AFM tip. It is our interpretation that the forces required to disrupt the complex on PS containing membranes are much lower, thus making it difficult to image. It is important to note that we cannot definitively differentiate whether the G-domain is bound to the bilayer or if it remains above the surface. In contact mode AFM the imaging forces result in height profiles that are typically much smaller than the actual height due to slight deformation of the sample..^28^ The results taken together with the rotational correlation times are fully consistent with the KRAS4b G-domain favoring a conformation that is in contact with the bilayer surface on PIP2 containing membranes.

### Binding KRAS4b to a mixed bilayer system

In previous studies we investigated the effect of relatively high mole fractions of anionic lipids on KRAS4b lipid interactions. In the plasma membrane PS and PIP2 constitute approximately 10% and 1% of the total lipid content respectively.^29,30^ Mixed lipid systems have not previously been investigated. Here we generate Nanodiscs containing 100% DMPC, 10% DMPS/90% DMPC and 10% DMPS/2.5% PIP2/87.5 % DMPC and measured the change in affinity (Figure 5). There is a striking increase in affinity in the presence of just 2.5% PIP2. Fitting the data to a simple Langmuir binding isotherm gives dissociation constants of 12 μM, 5.7 μM, and 1.4 μM for DMPC, 10% DMPS and 10% DMPS/2.5% PIP2 respectively. The presence of PIP2 increases the affinity 4-fold, which is striking considering that a 10 nanometer diameter Nanodisc has approximately 80 lipids/leaflet. This correlates to just two PIP2 per leaflet indicating that the interaction with PIP2 is favored over that of PS that is present at a concentration four times higher.

**Figure 5.**
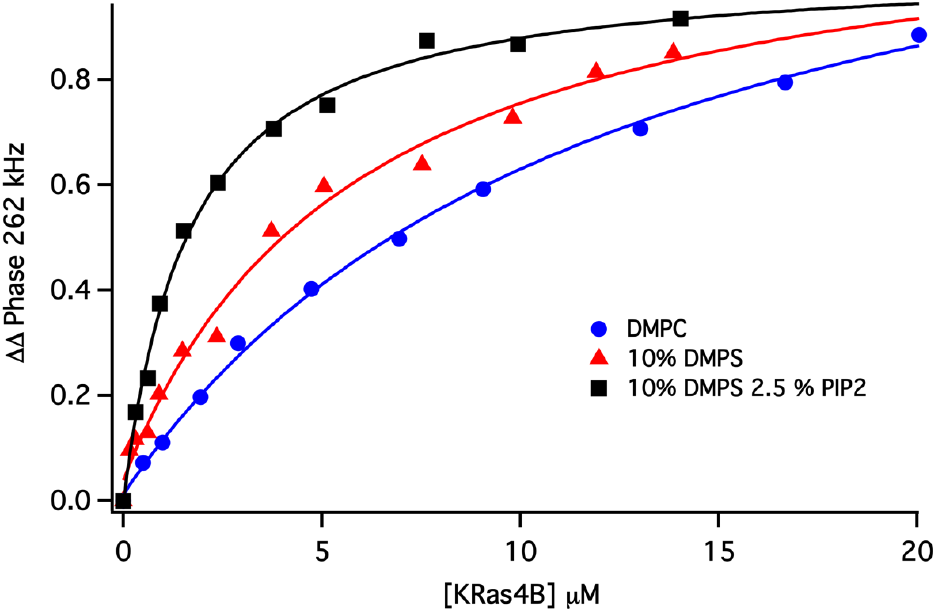
Single frequency KRAS4b binding isotherms. Blue 100% DMPC, Red 10% DMPS, Black 10% DMPS 2.5% PIP2. Solid lines represent fits to a simple Langmuir binding isotherm.

To gain a structural insight into the mechanism of KRAS4b – PIP2 interactions we performed multiple long term all atom MD simulations using a system comprised of a Highly Mobile Membrane Mimetic (HMMM)^31^. This membrane mimetic replaces the lipid tails with an organic solvent, dichloroethylene, allowing the lipid headgroups to diffuse rapidly and thus sample many lipid head group/protein interactions on a shorter time scale. Simulations one microsecond in length were performed on both 10% DMPS and 10% DMPS 3% PIP2. Starting configurations (five total) were generated from simulated annealing of the HVR (see methods, supplementary Figure S1) giving rise to 5 microseconds for each lipid system and 10 microseconds in total simulation time.

We first analyzed the trajectories by measuring the mean residence times for PIP2 and DMPS binding to each residue in the KRAS4b sequence. Interactions are defined as any lipid headgroup atom (excluding hydrogens) within 3.2 Å of a protein atom. Figure 6 shows the results for 10% DMPS (top panel) and 10% DMPS 3% PIP2 bottom panel. PS residence times vary from a few ns to a maximum of 16 ns over the combined simulation time (10 μs). Dramatically, many of the PIP2 residence times are significantly longer. Several Arg residues display residence times of more than 100 ns, indicating that the long-lived interactions are selective for Arg residues. Delocalized charges on the arginine headgroup and the multivalent charge associated with PIP2 give rise to these stable interactions. These results are in complete agreement with the increase in affinity shown in Figure 5.

**Figure 6.**
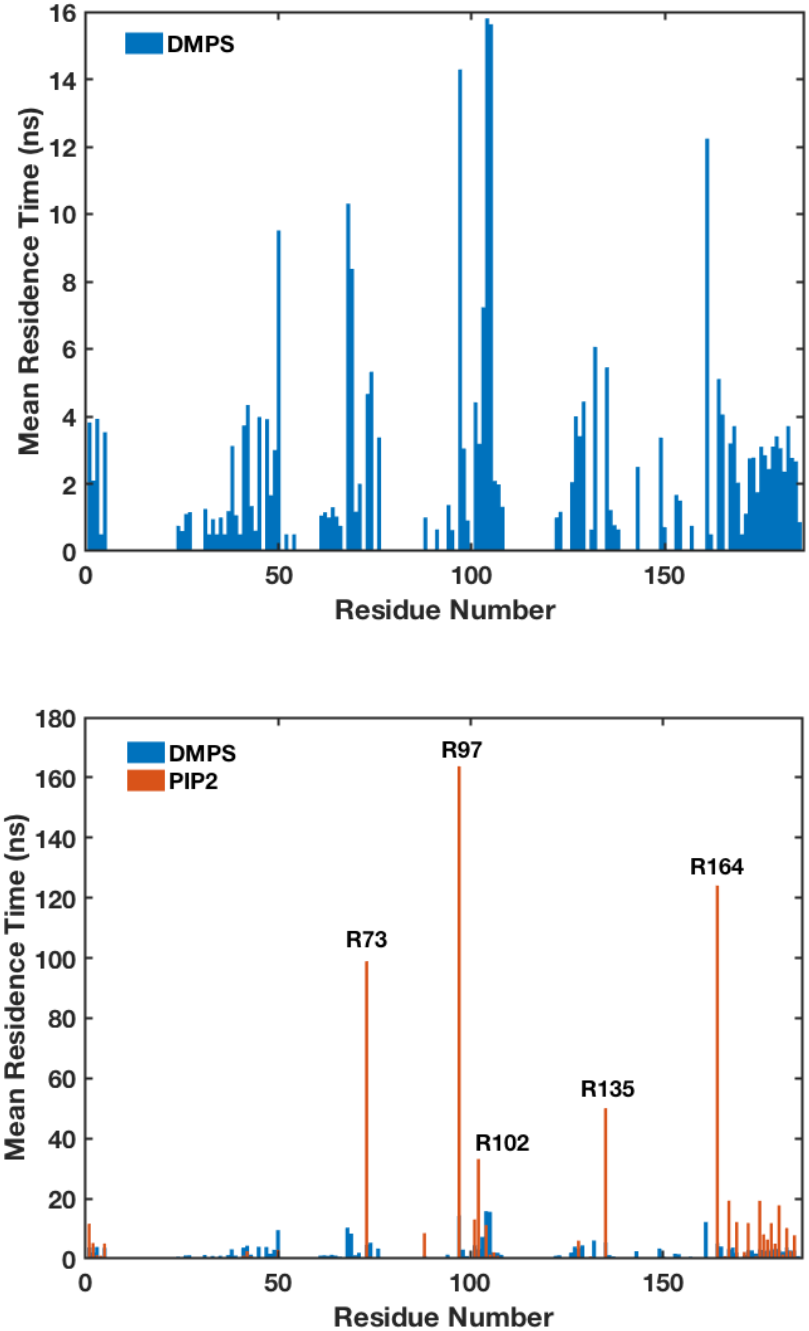
PIP2 Residue selectivity: Top panel: DMPS residence time. Bottom Panel:PIP2 residence times (orange), DMPS residence times are overlaid in blue. Note the scale change.

As a result of the long lived PIP2 protein interactions the conformational equilibrium is significantly changed. Figure 7 shows a two dimensional histogram of cos(θ) *vs*. distance for 10% DMPS (left) and 10% DMPS/3% PIP2 (right). A significant difference between the conformations sampled is observed. The black circles indicate the starting orientation for each of the 5 simulations. The red contour lines are the distribution fits and are summarized in Table 2. On bilayers containing only PS head-groups, the KRAS4b orientation redistributes significantly, sampling many configurations. The peaks in the distribution are far from the starting points. Simulations on bilayers containing PIP2, on the other hand, shows very tight distribution of orientations, and the distribution peaks remain near the starting points of the simulations. The narrow distributions are a consequence of the long-lived interactions between Arg residues and PIP2. These salt bridges do not allow for equilibration of the KRAS4b orientation on the timescales sampled, thereby suggesting that PIP2 could play a role in stabilizing KRAS4b orientation. We do not believe that PIP2 confers a specific orientational preference for KRAS4b but it stabilizes the KRAS4b-membrane complex and that the orientations sampled are a reflection of the initial starting configurations.

**Table 2.** KRAS4b Orientation Distributions

| KRAS4b-GDP 10% DMPS |  |  | KRAS4b-GDP 10% DMPS 2.5% PIP2 |  |  | G12V KRAS4b GTP 20% POPS <sup>9</sup> |  |  |
| --- | --- | --- | --- | --- | --- | --- | --- | --- |
| D(Å) | Cos($\theta$) | Orientation | D(Å) | Cos($\theta$) | Orientation | D(Å) | Cos($\theta$) | Orientation |
| 10.8 | 0.05 | Exposed | 9.7 | -0.37 | Exposed | 18.6 | -0.5 | OS1 |
| 28.6 | 0.02 | Occluded | 32.4 | 0.37 | Occluded | 33.3 | 0.9 | OS0 |
| 40.2 | 0.89 | Intermediate | 37.4 | 0.85 | Intermediate | 49.7 | 1.0 | OS2 |
|  |  |  | 37.4 | 0.54 | Occluded |  |  |  |

**Figure 7.**
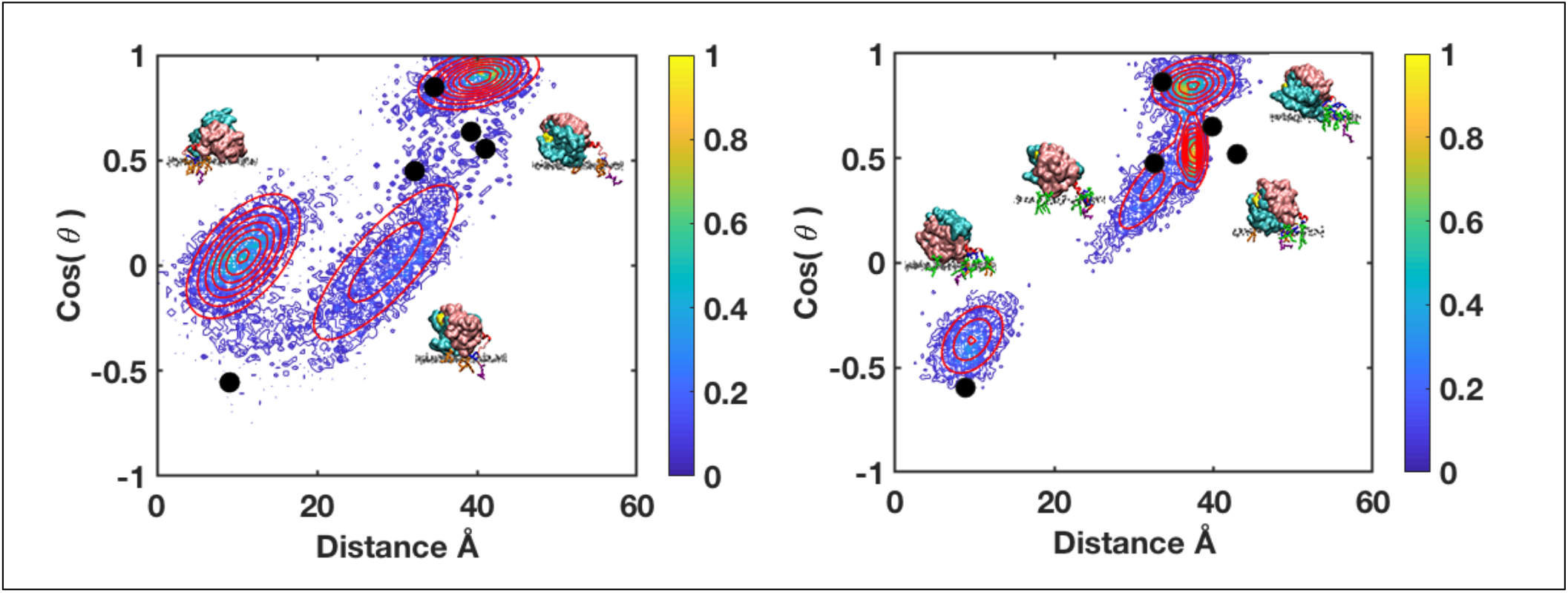
KRAS4b orientation. Left panel: cos (θ) *vs.* distance 10% DMPS. Right Panel: cos (θ) *vs.* distance 10% DMPS/3% PIP2. Black dots indicate starting orientations. Redlines are fits to a 2D gaussian distribution. Representative structures are for each distribution are shown. Pink=lobe 1, cyan = lobe2.

Interestingly if we compare our simulations of KRAS4b-GDP on DMPS to those of Prakash of KRAS4b G12V-GTP on 20% POPS, we see different orientations (Table 2).^9^ These authors show that KRAS4b-GTP prefers a configuration with an exposed effector binding site at cos(θ) = -0.5 and distance of 18.6 Å. In our simulations on 10% DMPS bilayers the orientation distributions differ slightly (Table 2). To define the functional role of the orientation of GDP bound KRAS4b we docked the GDP-GTP exchange factor SOS1 (pdb 4NYJ)^32^ on to the resulting structures by aligning the G-domain backbone of HRAS with KRAS4b in our simulations (Figure 8). In the docked structures we define exposed as orientations that allow interaction of SOS without interreference from the bilayer, intermediate as those with slight steric clash and occluded as those with significant overlap between the bilayer and SOS. Although the resulting orientations differ from that of KRAS4b-GTP on POPS membranes, the 3 resulting populations on 10% DMPS membranes can similarly be described as exposed, intermediate and occluded. These findings are consistent with the effect of GTP loading on the orientation of KRAS4b at the membrane surface^5,6^. On PIP2 containing bilayers of the 4 populations correspond to one exposed orientation, one intermediate orientation, and two occluded orientations. Thus, the majority of the orientations seen, exposed and intermediate, would allow for efficient interaction with the GDP-GTP exchange factor, SOS promoting activation of KRAS4b.

**Figure 8.**
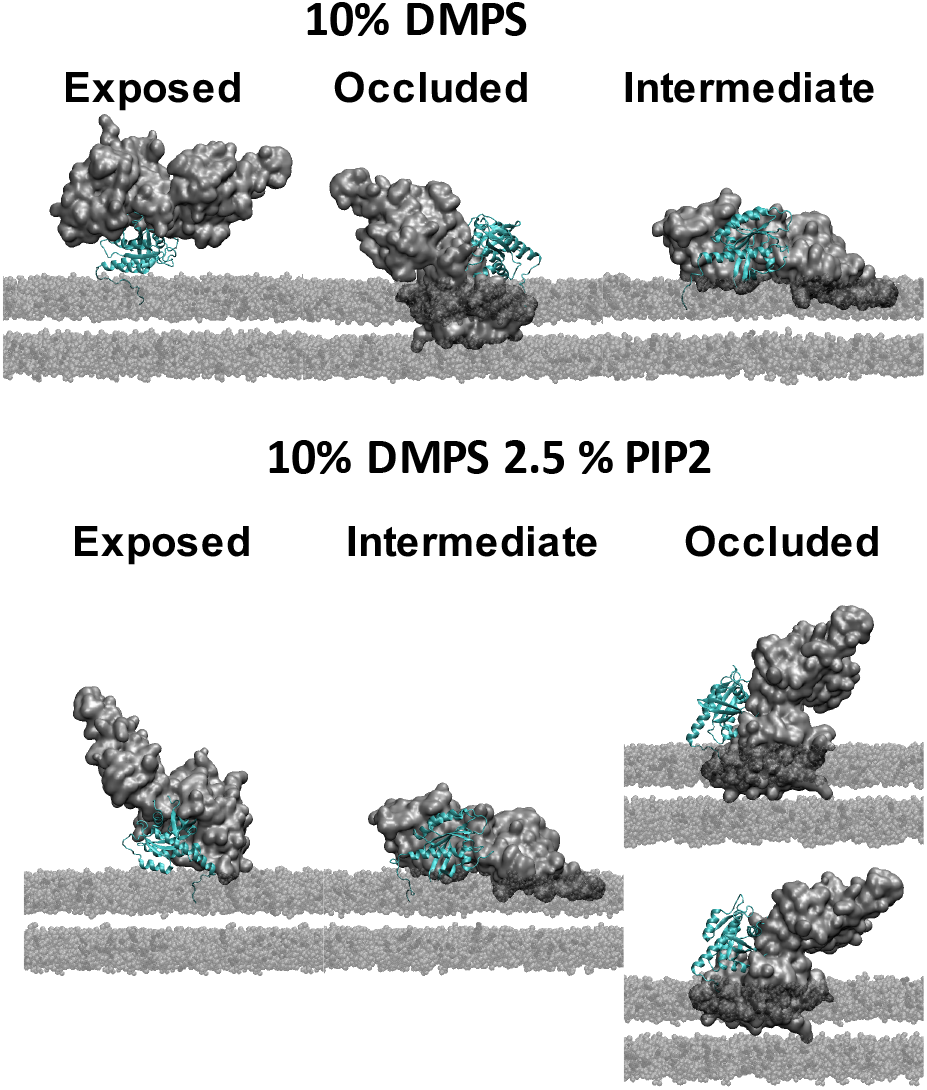
KRAS4b – SOS1 complexes. SOS1 (gray) docked onto the resulting orientations of KRAS4b (cyan). Top, 10% DMPS. Bottom 10% DMPS 2.5% PIP2.

An alternative way to view the effect of PIP2 on KRAS4b membrane dynamics is to look at the mobility of the G-domain when tethered to the bilayer surface. The mobility is a measure of how fast the orientation is changing at any point during the trajectory. The difference between KRAS4b on DMPS *vs*. DMPS/PIP2 membranes can be visualized in a two-dimensional histogram of the mobility of each orientation *vs.* the angle. Here again we see that the distribution of the mobility over orientation angle is quite broad on 10% DMPS (figure 9, top), while on 10% DMPS/3% PIP2, the mobility is much smaller and displays a narrow distribution (figure 9, bottom). Our interpretation is that just a small amount of PIP2 will slow the exchange of KRAS4b from the exposed OS1 state to the occluded OS2 state. This only reflects a stabilization of the orientation not a selective preference of one orientation over the other. What is intriguing is the fact that just a few PIP2 will promote this stability. Therefore, the colocalization of PIP2 could dramatically influence the orientation of KRAS4b and thereby its interaction with PI3K or the nucleotide exchange factor SOS resulting in altered KRAS4b activity and PI3K signaling.

**Figure 9.**
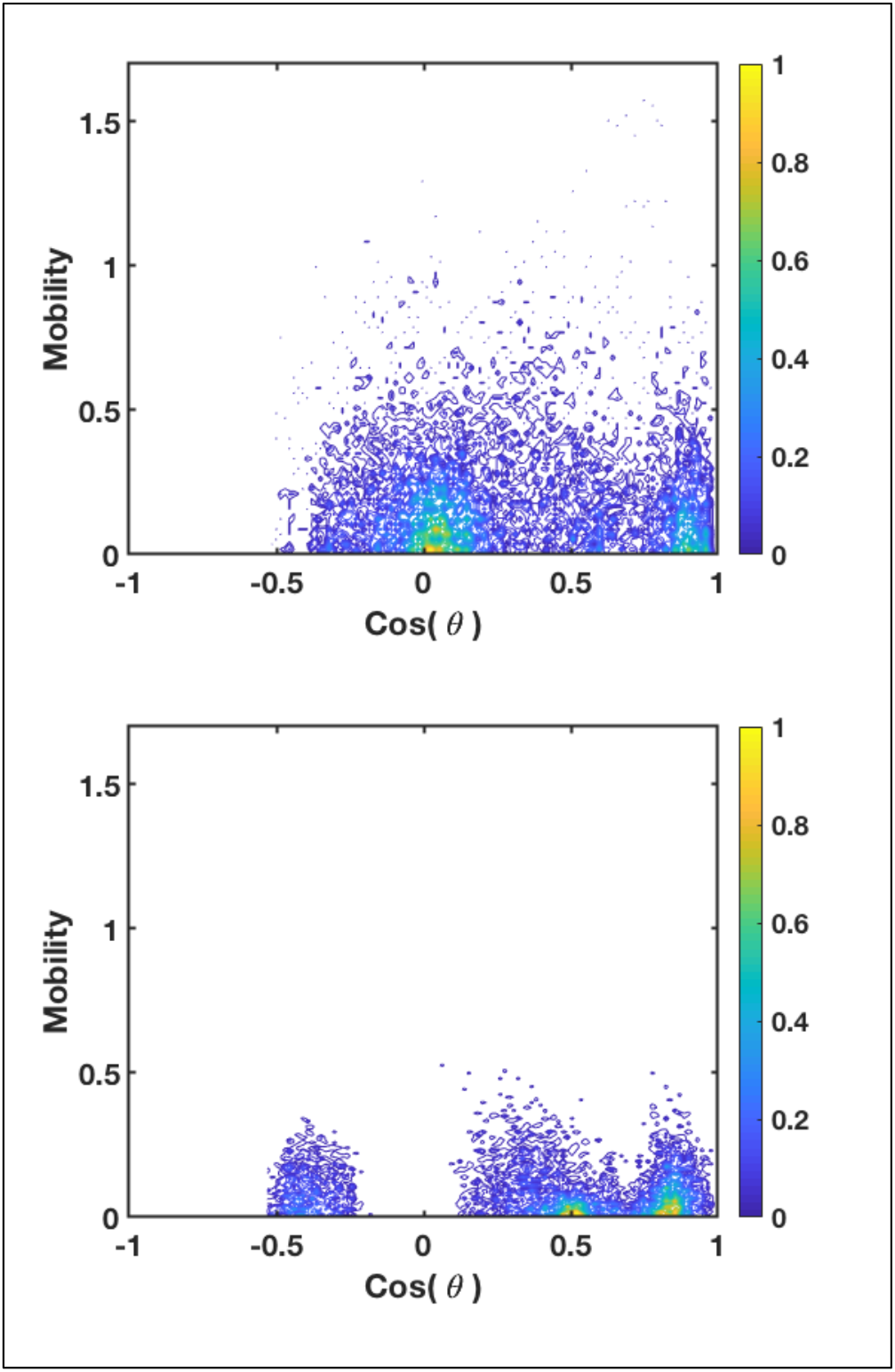
KRAS4b Mobility. Top panel: Mobility *vs*. cos (θ) DMPS. Bottom Panel: Mobility *vs*. cos (θ) 10% DMPS/3% PIP2

## CONCLUSIONS

Using experimental and theoretical analysis we present a picture of KRAS4b interaction with anionic lipids. On PS containing membranes the KRAS4b-GDP samples many configurations and is loosely associated with the bilayer surface. KRAS4b is dynamic on PS membranes favoring a more occluded state, consistent with the inactive GDP bound form. However, on membranes containing PIP2, the G-domain is tightly associated with the bilayer surface, and transitions between the orientations are much less frequent. We propose that PIP2 stabilizes G-domain bilayer interactions, regardless of orientation as long as there are accessible Arg residues. Owing to the importance of PIP2 as a signaling lipid and its role in downstream effects, the stabilization of the supramolecular complexes by PIP2 could play a role in RAS dependent signaling.

## ASSOCIATED CONTENT

Figure S1 generation of MD starting configurations.

Figure S2 Definition of orientation parameters and orientational mobility.

## AUTHOR INFORMATION

### Author Contributions

M.A.M designed and carried out the experiments and wrote the manuscript. A.G.S provided KRAS4b samples and contributed to experimental design and writing of the manuscript. S.G.S is the project leader and contributed to experimental design and writing of the manuscript.

## ACKNOWLEDGEMENTS

We thank Peter Frank, Kelly Snead, Dominic Esposito, William Gillette and Jennifer Mehalko from the NCI-RAS Initiative for production of farnesylated and methylated KRAS4b protein used in this work.

## ABBREVIATIONS

PIP2: L-α-phosphatidylinositol-4,5-bisphosphate
DMPC: 1,2-dimyristoyl-sn-glycero-3-phosphocholine
DMPS: 1,2-dimyristoyl-sn-glycero-3-phospho-L-serine
HVR: hyper-variable region
HMMM: Highly Mobile Membrane Mimetic
PM: plasma membrane
ND: Nanodisc

