## Supplementary material for "PIP2 Influences the Conformational Dynamics of Membrane bound KRAS4b": Supplemntal information

Supplemental Information

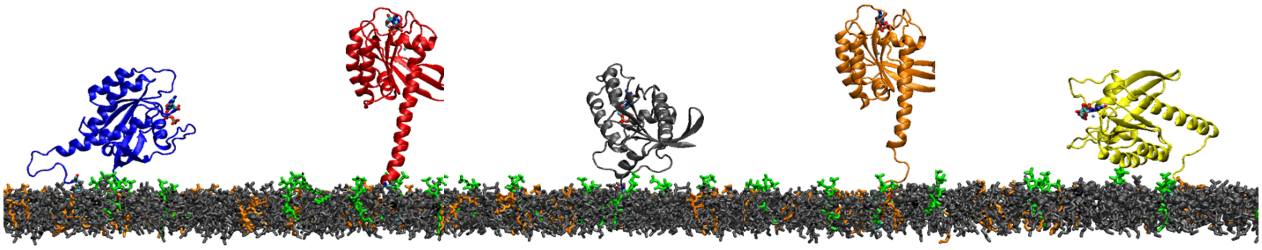

Figure S1. Five separate simulated annealing molecular dynamic runs of the KRAS4b HVR were performed by heating the system to 50 C over 50 ns followed by cooling to 20 C over 50 ns. The G-domain was constrained during the annealing steps. A final equilibration of 100 ns was performed at 20 C to relax the entire system. The 5 resulting structures were docked onto the membrane using CHARMM-GUI HMMM builder.

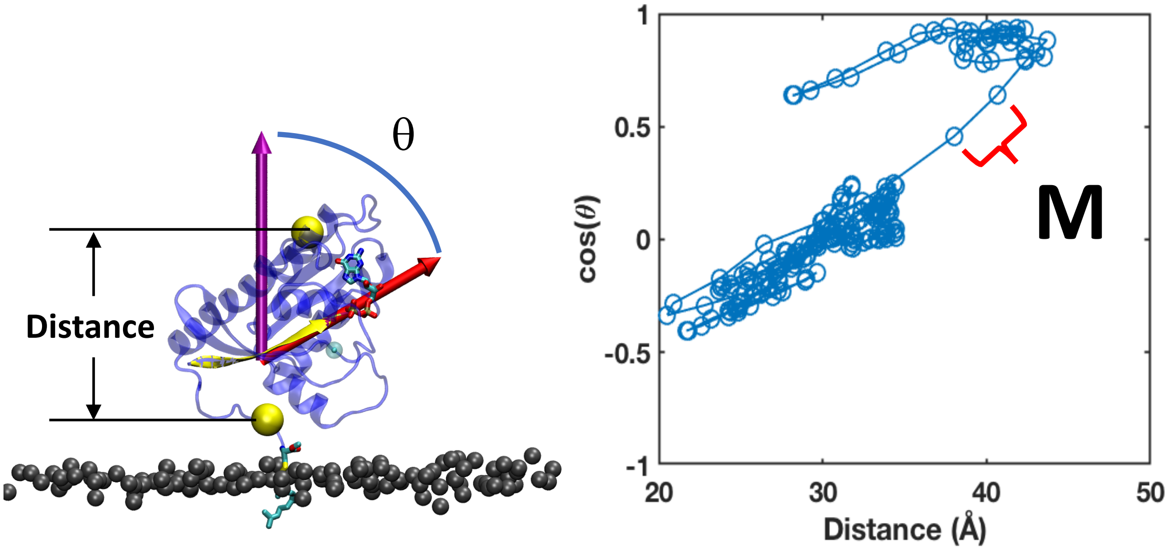


Figure S2 Orientational Mobility

The orientation is defined by the angle, $\theta$, of $\beta$-sheet 1 with respect to the bilayer normal and the distance of $\alpha$-helix 4 above the bilayer surface. The Mobility is defined as the distance between consecutive points (500 ps / pt) on the orientation plot of cos($\theta$) vs Distance, giving a measure of the change in orientation per time.
